# Actin bundles play different role in shaping scale as compare to bristle in mosquito *Aedes aegypti*

**DOI:** 10.1101/2020.04.06.027110

**Authors:** Sanja Djokic, Bakhrat Anna, Ido Zurim, Nadya Urakova, Jason L. Rasgon, Uri Abdu

## Abstract

Insect epithelial cells contain cellular extensions such as bristles, hairs and scales. It has been suggested that these cellular extensions are homologous structures that differ in morphology and function. These cellular extensions contain actin bundles that dictate their cellular morphology; bristle and hair are cylindrical in shape, while scales are wider and flattened. While the organization, function and identity of the major actin bundling protein in bristles and hairs is known, this information in scales is unknown. In this study, we characterized the development of scales and the role of actin bundles in the mosquito, *Aedes aegypti*. We show that scales undergo drastic morphological changes during development, from cylindrical shape to flat shape with longer membrane invagination. Scale actin bundle distribution changes during development, from symmetrical organization of actin bundles located throughout the bristle membrane, to asymmetrical organization of the actin bundles. By chemically inhibiting actin polymerization and by knocking-out the *forked* gene in the mosquito (*Ae-Forked;* a known actin bundling protein), by CRISPR-Cas9 gene editing, we showed that actin bundles are required for shaping bristle, hair and scale morphology. We demonstrated that actin bundles and *Ae-Forked* are required for bristle elongation, but not that of scales. In scales, actin bundles are required for width formation. Our results reveal a differential requirement of actin bundles in shaping mosquito scales compared to bristles.

## Introduction

The vast majority of the epithelial cells that cover insects ‘contain cellular extensions; namely bristles, hairs and scales. In *Drosophila*, it has been suggested that thoracic bristle, wing hair, and scales are homologous structures that differ in their morphology (Galant et al., 1998, Guild et al., 2005, Zhou et al., 2009). Bristles are elongated and cylindrical in shape, with a different morphology and function than the hairs, which are short and shaped like rose thorns, and scales, which are flattened and thin.

Most of our knowledge on the role of actin in bristle (Tilney and DeRosier, 2005) and hair (Mitchell et al., 1983; Turner and Adler, 1998; Guild et al., 2005; Ren et al., 2006) development are from studies on *Drosophila*. The bristle contains membrane-associated actin filament bundles (Tilney et al., 2000a; 2000b). It has been shown that two actin cross-linker proteins are involved in bristle actin bundle formation; the first is Forked, the human epsin protein homologue (Tilney et al., 1995; Tilney et al., 1996). The second is Singed, the *Drosophila* Fascin homologue; hereafter, referred to in the text as Fascin, (Tilney et al., 1995; Paterson and O’Hare, 1991; Bryan et al., 1993). In hair, there are no membrane associated actin bundles but instead overlapping cytoplasmic bundles. Organization of actin bundles in the hair is achieved by the sequential use of three actin bundling proteins: Villin, Forked and Fascin. Thus, bristle and hair generate actin filament bundles but employ different strategies to assemble these into vastly different shapes (Guild et al., 2005). There is not much information on the role actin bundles in shaping insect scales (Dinwiddie et al., 2014; Day et al., 2019). Recently, it was shown that in the butterfly, *Vanessa cardui*, actin bundles in the scales were required for initial scale elongation and to orient the scale parallel to the wing membrane. On the morphological level, actin bundles are required for longitudinal ridge formation and for producing finger-like projections at the tips of wing scales (Dinwiddie, et al 2014).

Besides *Lepidoptera*, it is also known that in several mosquito species, the entire body is covered with scales. It was suggested that leg scales may have a role in egg laying by giving the mosquito high water buoyancy and floating ability (Wu et al., 2007). In this study, we show that during *Aedes aegypi*, pupal development, scales undergo drastic morphological changes. In early pupal stages the scales are cylindrical in shape, then flatten with longer membrane invagination. Actin bundle distribution is changed during development, from symmetrical organization of actin bundles located throughout the bristle membrane, to asymmetrical organization of round vs. flattened actin bundles. To study the role of actin bundles in scale development, we inhibited actin polymerization during pupal development using chemical inhibitors, and used CRISPR-Cas9 gene editing to knock out the mosquito actin bundling gene *forked* (*Ae-forked*). We found using both methods that, scale morphology was altered in ridge organization and shape. We also determined differences in the role of actin bundles in cell elongation of mosquito scales to bristles vs. hairs. Our results revealed that the unique organization of scale actin bundles dictate their cellular morphology.

## Results

To understand the role of actin on mosquito scale development, we focused our analysis on four different body sections, namely, the thorax (Fig. 1 A, A’), scutellum (Fig. 1 B, B’), abdomen (Fig. 1 C, C’) and legs (Fig. 1 D, D’). The mosquito thorax is covered with falcate scales (Fig. 1 A, A’), which are curved with a sharp or narrowly rounded apex. At the scutellum, we found that each of the three scutellar lobes (Fig. 1 contains clusters of five bristles each with its associated spatulate scales, which are lamellar characterized by a broad distal section and attenuation at the base (Figure 1 B’). Also, the abdomen (Fig. 1 C) and the legs (Fig. 1 D) are covered with spatulate scales (Fig. 1 C’, D’). As previously described (Wu et al., 2007), along each scale, there are longitudinal ridges spreading from the scale base to tip (Fig 1 A’, B’,C’, D’). Closer examination revealed that only thorax (Fig. 1A’’), abdomen (Fig. 1C’’) and legs (Fig. 1 D’’) but not scutellum (Fig. 1 B’’) scales had cross ribs between the ridges.

**Figure 1:**
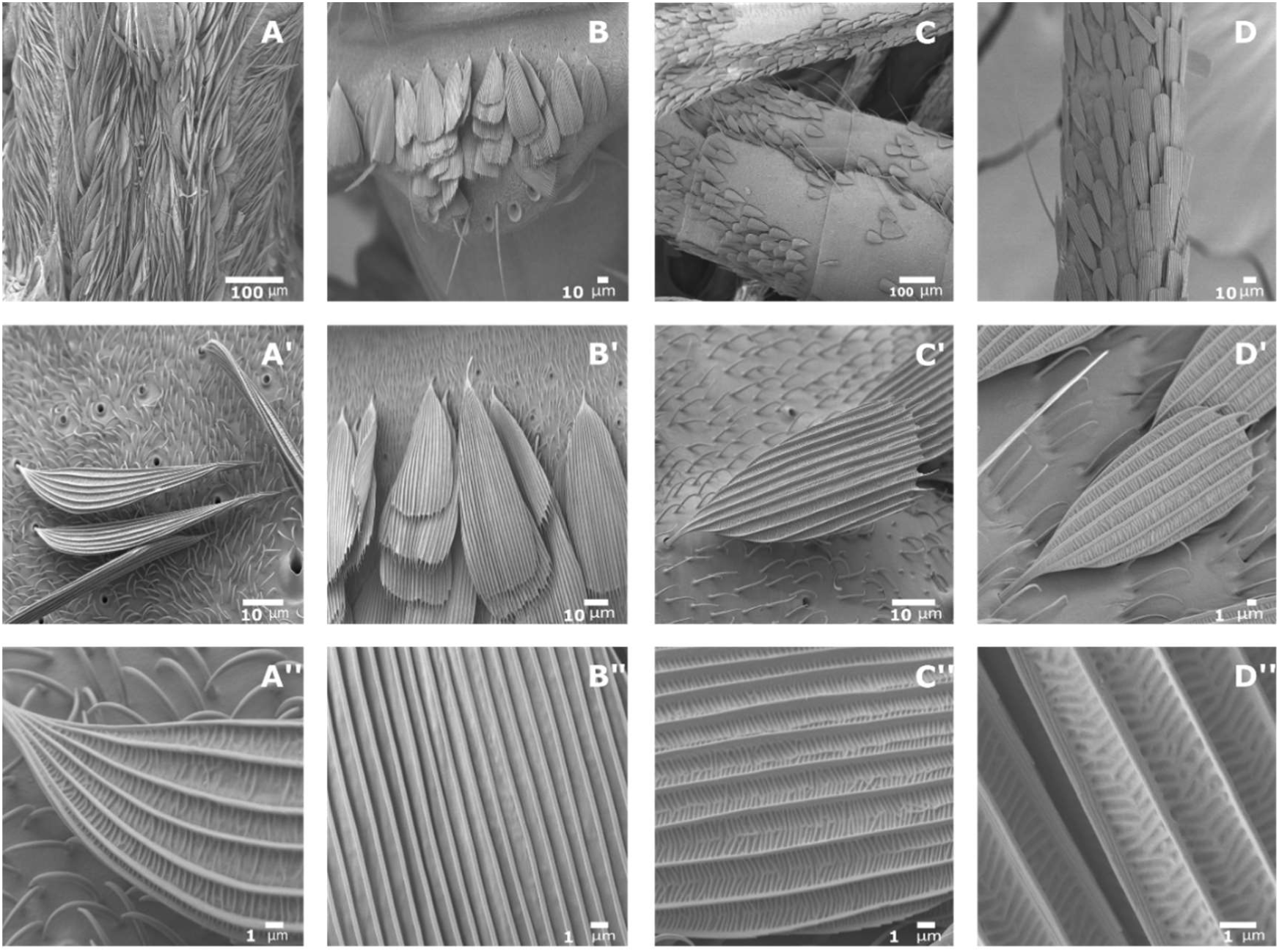
Scanning electron microscopy (SEM) images of scale from different tissues of the mosquito *Aedes aegypti*. A) Thorax, B) Scutellum, C) Abdomen D) leg, from mosquito. (B’) Closer examination of the scales reveals that scales on the thorax (A’) are falcate type, and the one on the Scutellum (B’) Abdomen (C’) and leg (D’) are spatulate type. A long each scale, there are longitudinal ridges which spread from the base of the scale to the tip (A’’-D’’). Exept from scale on Scutellum (B’’) in all other scales between the longitudinal ridges, there are many cross ribs. Scale bars are found on each image.

Next, using actin staining, we followed both legs and scutellum scale bud initiation (Fig. 2). We found that on the scutellum, bristles emerged as early as 5 hours after pupa formation (APF) (Fig 2 A), but buds of scales were detected only at 7 hours APF (Fig. 2B). On the legs, scales arise as little buds as early as 5 hours (Fig 2 APF. Scales on the legs were found in clusters of four (Fig. 2 D), and in each cluster, the scales differed in size (Fig. 2 C’, D’). Our results demonstrate a difference in timing of scale development between body sections, as well as a difference in scale and bristle development on the same section.

**Figure 2:**
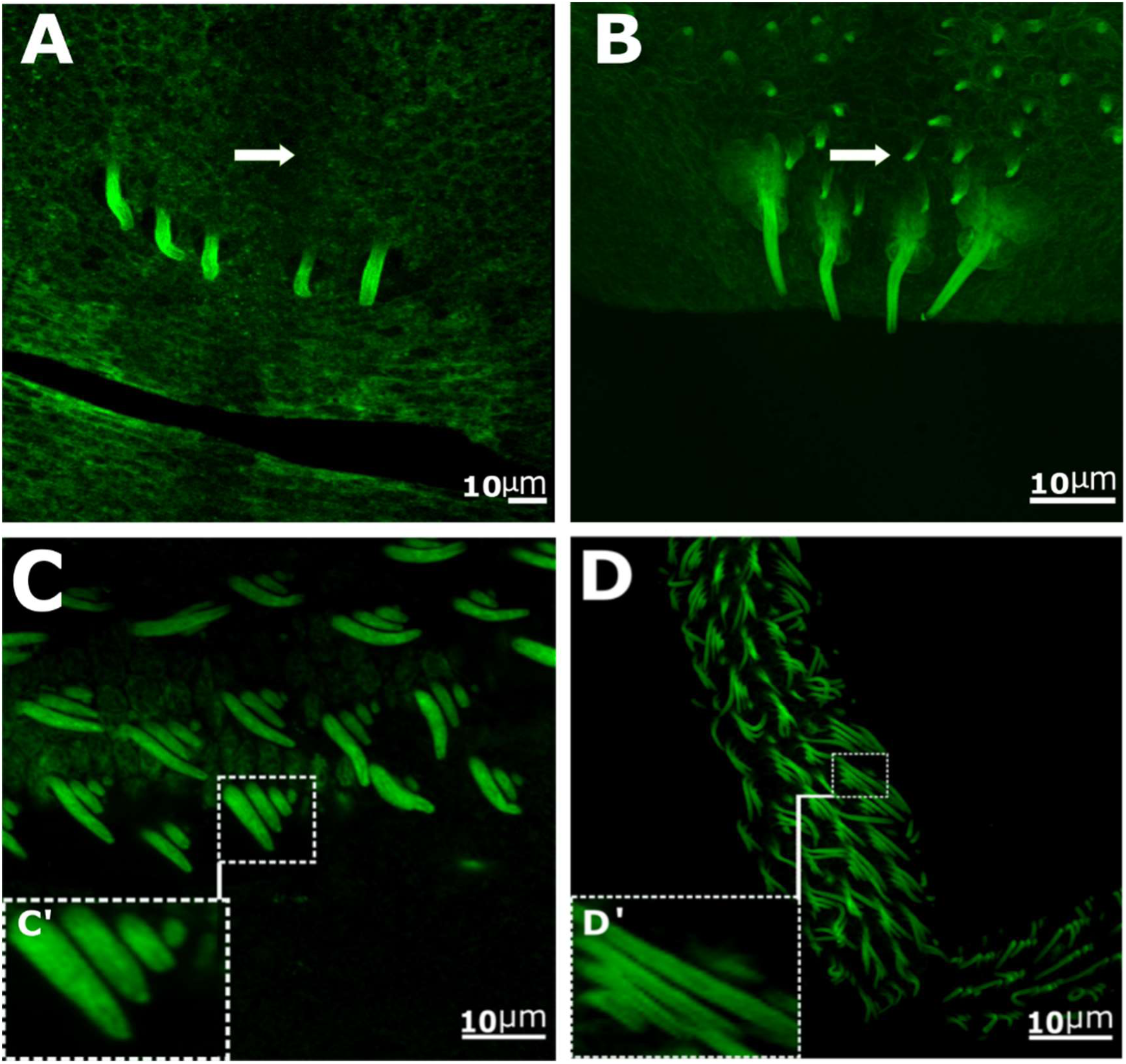
Confocal images of mosquito scale development. A-D, phalloidin staining (actin-green) of Scutellum (A-B) and leg (C-D) from mosquito pupa at different developmental time. Scutellum of pupa 5h (A) and 7h (B) after pupa formation (APF), at 5h small buds of bristles can be detected (arrows), but not scales. At 7h (B) both bristle and scale (B’) could be detected. Leg of pupa 5h, (C) and 7h APF (D), scales in clusters of four can be seen (C’, D’).

### Ultrastructure analysis of the mosquito scale

To follow ultrastructure changes of leg scale development, we used TEM analysis. We chose to use legs for EM analysis since this body part is easier to handle compared to the thorax. At 7 APF, scales were cylinder in shape, resembling bristle, with membrane-associated actin bundles and dense microtubules (MTs) found distributed over the scale cytoplasm. The scale contained about 20 rounded actin bundles that were similar in size throughout the scale membrane (Fig. 3 A–C). At 10h APF (Fig. 3 D–G), scales were still rounded with a dense MT network all over them, but this time, the actin bundles had grown in size and were triangular in shape. The bundles were organized asymmetrically with one side of the scale containing large triangular bundles, with much smaller rounded ones on the opposite side. At 12h APF (Fig. 3 H–K), the scales became flattened with now centrally localized MTs. Large rounded actin bundles were on one side, and there were smaller and flatter ones on the opposite side. At 18h APF (Fig. 3 L–N), the flattened scales had larger ridges on one side with smaller, flatter actin bundles, and on the other side, rounded larger bundles could be seen. At this stage, only a few MT filaments were visible.

**Figure 3:**
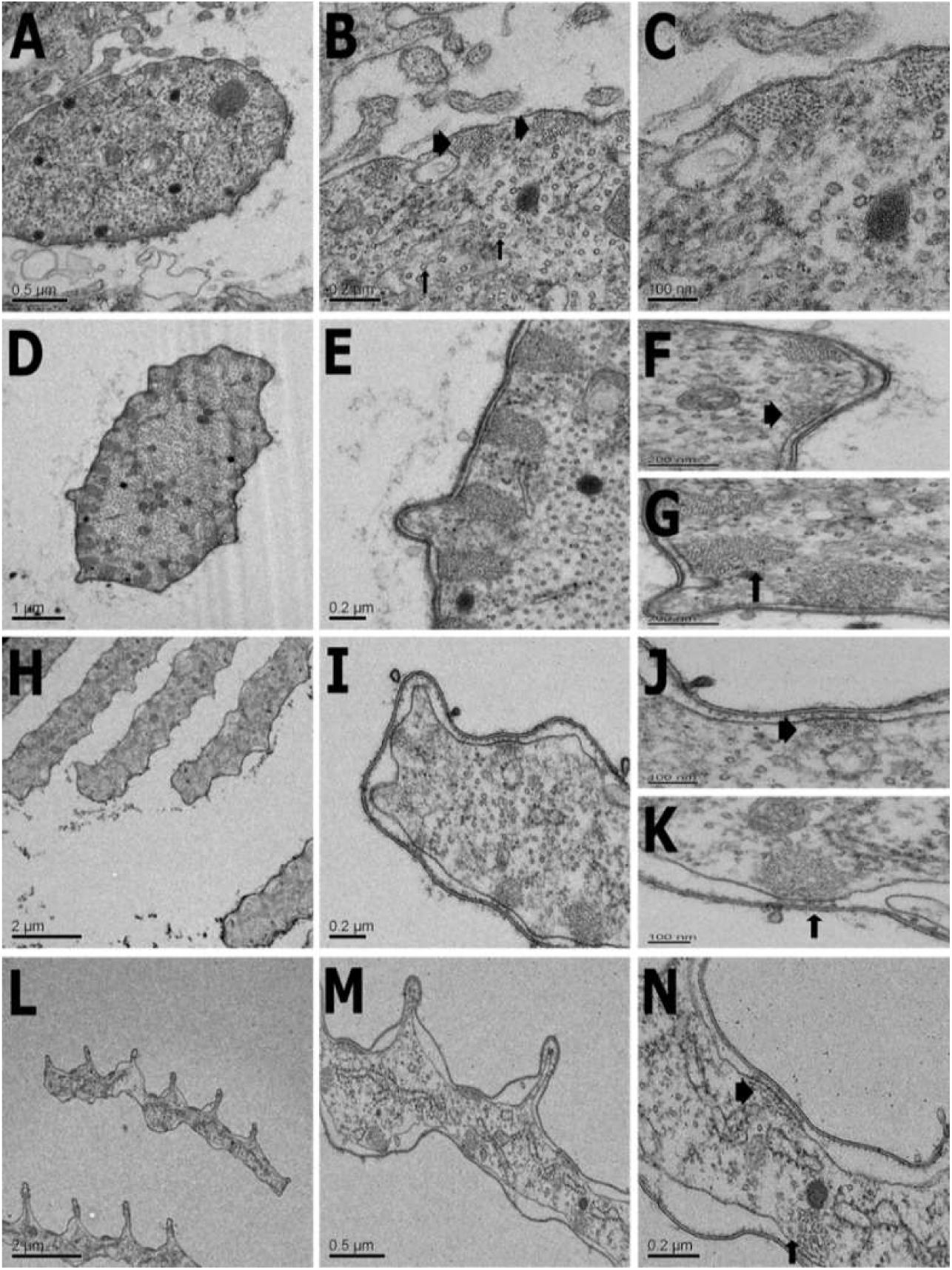
Transverse transmission electron microscopy section of *Aedes aegypti* scales from the leg during development. (A–C) Scales at 7h after pupal formation (APF) show (B) centrally positioned microtubules (arrows) and actin bundles (arrowhead) attached to a plasma membrane. (D–G) At 10h APF, actin bundles are organized asymmetrically. (G) One side of the scale contains large triangular actin bundles (arrows), (F) and on the opposite side, much smaller actin bundles are evident (arrowhead). (H–K) 12h APF scales are flattened and elongated, and actin bundles are organized in an asymmetrical manner. (J) Actin bundles are smaller on one side of the scale (arrowhead), (K) and much larger on the other side of the same scale (arrows). (L–N) Scales at 18h APF, one side of the flattened scales contains obvious ridges and valleys. These are on the side with flattened actin bundles. (N) At this stage, actin bundles have the same pattern with smaller scales on one side (arrowhead) and larger scales on the other (arrows). Scale bars are found on each image.

### Inhibition of actin polymerization by Cytochalasin D injection affect bristle and scale development

To study the role of actin in bristle and scale development, Cytochalasin D (CD) (which inhibits actin polymerization by capping the barbed end of actin filaments and inhibits further elongation from the barbed end [Cooper, 1987] was injected into pupae. Previously it was shown, that injection of CD into pupa affect *Drosophila* bristle growth (Fei et al., 2002; Guild et al., 2002). We focused analysis on scutellum bristles and scales. Since we found that on the scutellum initiation of bristles and scales start after 5 and 7 hours, respectively (Fig. 2), we injected CD into 4h AFP pupae. We found that the bristle length (148.791±8.803 µm) of treated pupae (Fig. 4 B) was significantly shorter (t=-7.101, df=6, p=0.0004) then wild type bristles (Fig. 4 A; 255.082±12.106 µm; Table 1). The overall morphology of the bristles were affected, instead of long tapered cylinder shape (Fig. 4C) with regular ridges and valley pattern (Fig. 4 C) the bristles were blunt with splits along their length (Fig 4 D) and the ridge and valley pattern was mis-orientated (Fig 4 C). Next, we examined scale morphology and found clear defects in their morphology. Overall, the scales were curved in shape (Fig. 4 B, G), and their length was significantly (t=3.156, df=13, p<<0.0076) longer (77.27±5.195 µm) than wild type (56.789±3.891 µm; Table 1). Moreover, the scales had lost their spatulate shape (Fig. 4B vs. 4D) where the upper part was no longer broad, but instead tapered towards the tip, resembling the bristle tip. They were significantly narrower (t=-11.385, df=12, p<<0.0001) then wild type, both in the middle; 7.029±1.072 µm vs. wild type=25.385±1.204 µm and (t=-23.406, df=5, p<<0.0001) at their tip; 1.09±0.111 µm as compared to wild type=23.718±0.960 µm; Table 1. Closer examination of the scale revealed that the longitudinal ridges were mis-oriented (Fig 4H).

**Table 1.** Bristle and scale measurement in WT and Cytochalasin-D treated pupa.

|  |  |  |
| --- | --- | --- |
| Genotype | WT | Cytochalasin-D |
| No. of pupa | 4 | 9 |
| No. of bristles | 15 | 30 |
| Bristle length | 255.082±12.106* | 148.791±8.803 |
| No. of pupa | 6 | 9 |
| No. of scales | 21 | 34 |
| Scale width- middle | 25.385±1.204* | 7.029±1.072 |
| Scale width-top | 23.718±0.960* | 1.09±0.111 |
| Scale length | 56.789±3.891* | 77.27±5.195 |

\* - represents a significant difference (P<0.05) between genotypes (for a detailed description of statistical analysis performed, see Materials and Methods).

**Figure 4:**
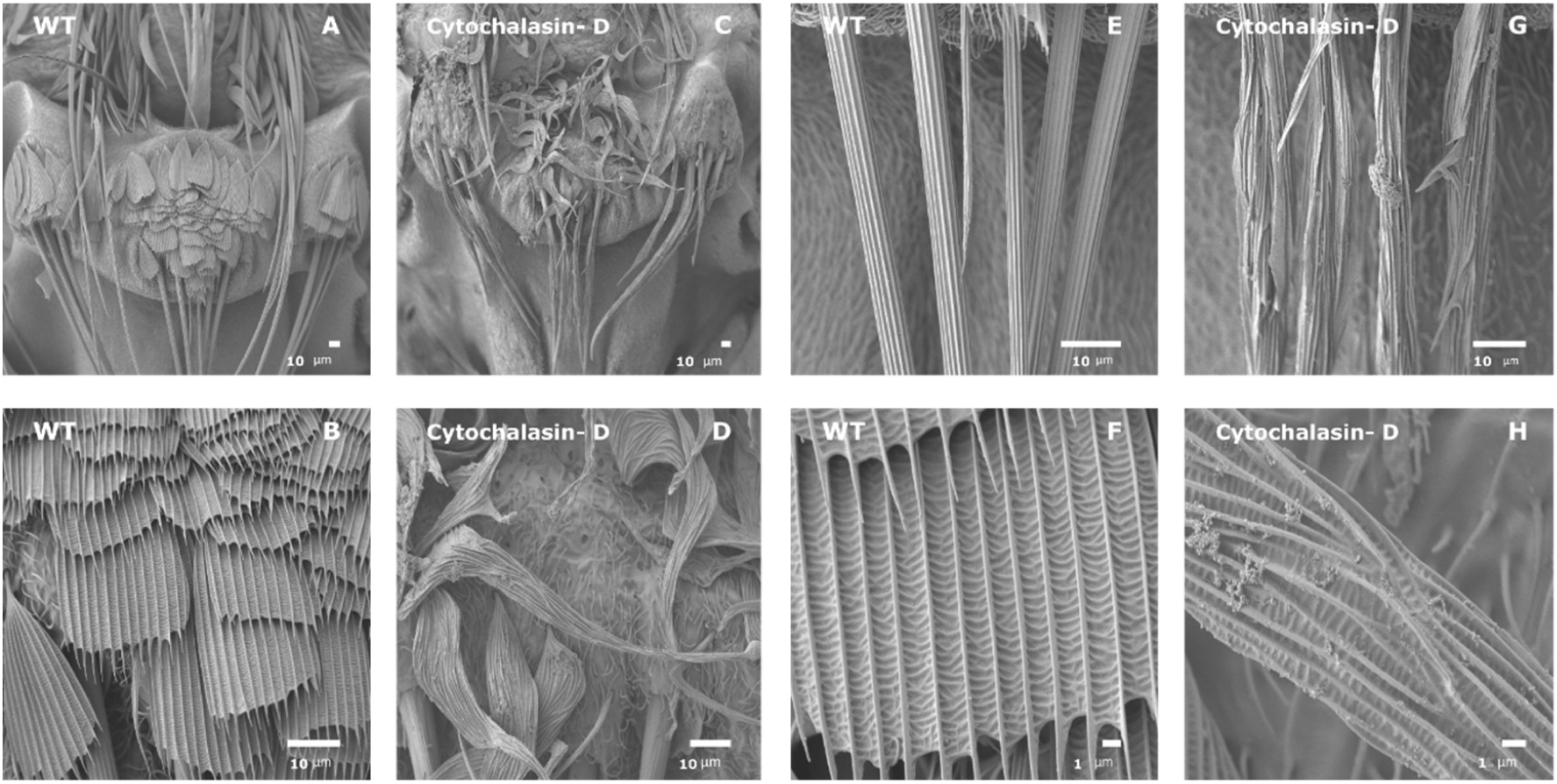
Scanning electron microscopy (SEM) images of Scutellum region from of WT and from cytochalasin-D (CD) treated mosquito *Aedes aegypti*. Scutellum region from WT (A) and CD-treated (C) mosquitos. Overall, the bristle are much shorter and scales are twisted in CD-treated mosquitos. Closer examination of bristle from CD-treated mosquitos (G) revealed mis-orientaion of the ridges pattern as compared to WT (E). Closer examination of scales from CD-treated mosquitos (D) revealed that the WT-like spatulate shape (B) is lost in CD-treated mosquitos and now is twisted, narrower and the upper part was no longer broad, but instead tapered towards the distal part of the scales. Also, mis-orientaion of the ridges pattern in CD-treated (H) mosquitos as compared to WT (F) is detected. Scale bars are found on each image.

### Forked is required for both mosquito bristle and scale development

To further investigate role of actin bundles in scale development, we mutated one of the known *Drosophila* bristle and hair actin bundles genes, (*forked*). Blast analysis revealed the annotated gene AAEL018112 (hereafter *Aa-forked*), was homologous to the *Drosophila forked* gene. To generate the *Aa-forked* mutant line, a sgRNA targeting exon 7 (Fig. 5A) was injected into eggs of a transgenic mosquito line that expressed Cas9 under control of the ubiquitin L40 promoter (Li et al., 2017). Injection of sgRNA into 200 embryos yielded 26 Generation 0 (G0) mosquitoes. All of the G0 mosquitoes were crossed to wild-type mosquitoes of the opposite sex. Next, eggs from one of the G1 generation lines were hatched and all 50 offspring were PCR-screened for detection of cas9-generated mutations. Among the 50 screened adults, 47 had 1 visible PCR product of the expected size of the wild-type allele (Fig. 5 B1), 3 individuals had two PCR products (one with the expected size of the wild-type allele, and a second smaller PCR product indicating a deletion [arrow in Fig. 5B]),DNA sequencing of the smaller band revealed a 52 bp deletion (Fig. 5) at the sgRNA site in *Aa-forked* gene, resulting in a predicted frameshift leading to a premature stop codon and a predicted truncated protein of 801 aa instead of the 1342 aa in the wild-type protein (Fig. 5A). We mated G2 heterozygous mosquitos and screened by PCR for homozygous *Aa-forked* mutant individuals. We found that *Aa-forked* gene is essential for mosquito development as all identified homozygous *Aa-forked* mutants (Fig 5 B3) died as pharate adults that were partly emerged from their pupal case.

**Fig. 5.**
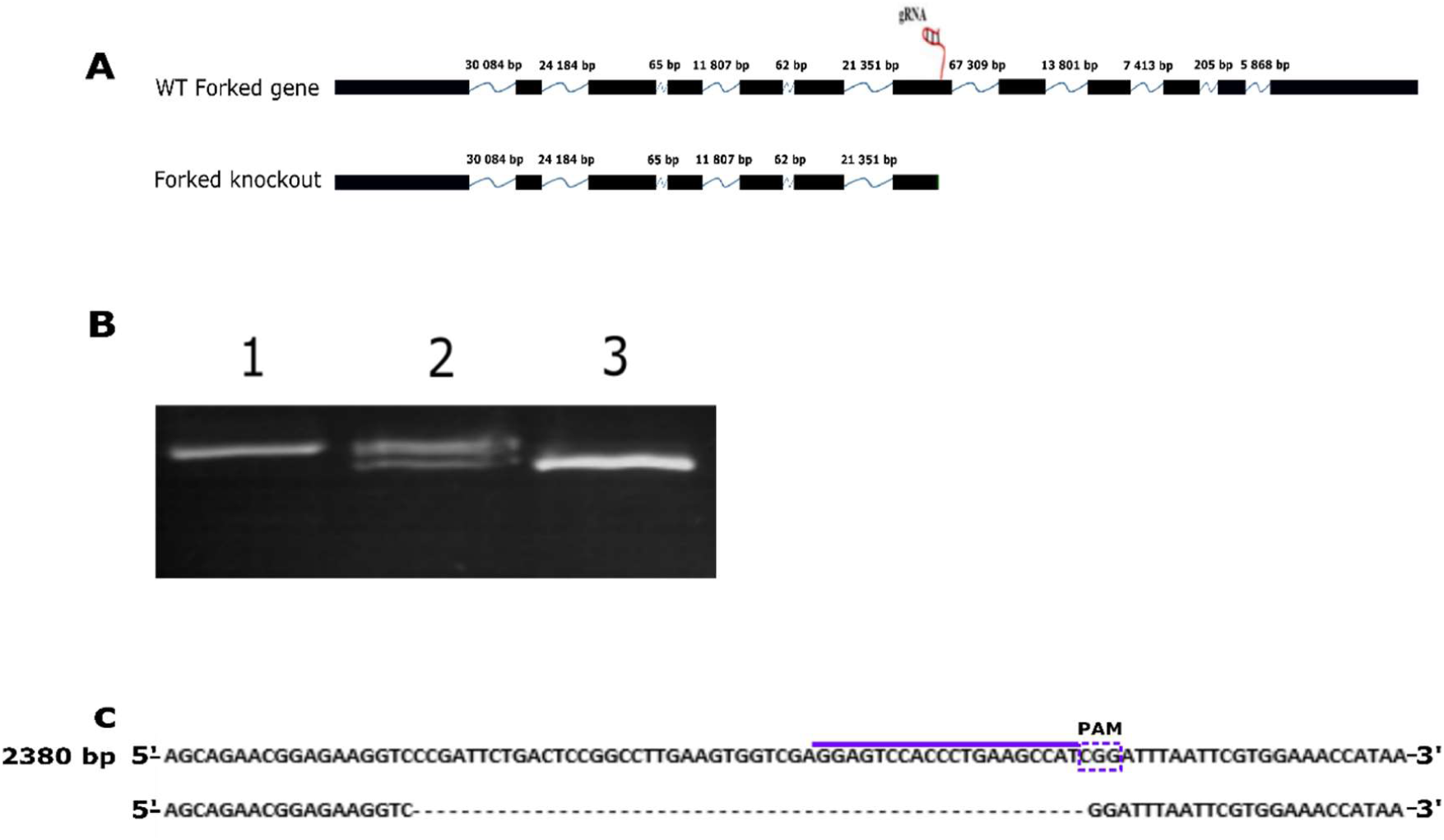
Generation *Ae-forked* mutant using CRISPR/CAS9 system. **A)** Genomic organization of *Ae-forked* gene, black boxes are exons, blue line represent introns, number above them is intron size in bas pairs. sgRNA on exon 7 is marked with red. sgRNA was injected to *A. aegypti* transgenic mosquitoes embryo ubiquity expressing CAS9 protein under *ubiquitin L40 (*AAEL006511) promoter were used (Li et al., 2017). **(B)** PCR analysis of WT and mutant mosquito lines. 1) PCR on WT mosquitos reveal PCR product of in size. 2) PCR on G1 putative mutant heterozygous lines reveal 2 PCR products, one in the same size as in WT the second one is smaller in size. 3) PCR on homozygous G2 *Ae-forked* line showing only one band which represent the deletion of 52 bp. **(C)** DNA sequence of genomic DNA from WT *Ae-forked* around the sgRNA site (upper line in purple end the PAM site is marked with box). Below is DNA sequence of genomic DNA from *Ae-forked* mutant line, showing deletion of 52 bp.

Next, we used SEM to examine the nature of the defects in both the hairs, bristles and scales in Aa-forked mutant lines. Examination by SEM (Fig. 6) showed that scales were affected in on the thorax (6 A-B) notum (Fig. 6, C-D) leg (Fig. 6, E-F) and abdomen (Fig. 6, G-H). In mutant mosquito body parts that had scales with spatulate shape such as the notum (Fig 6C) leg (6E) and abdomen (6G), the upper part was no longer broad as in the wild-type, but instead tapered towards the tip (Fig. 6 D, F, E). Examination of scales in all body parts revealed that, compared to wild type which had longitudinal ridges (Fig. A’, C’, E’, G’), in *Aa-forked* mutants all of the ridges were mis-oriented (Fig. B’, D’, F’, H’). To quantify the morphological changes in scales morphology in Aa-forked mutants, we focused our analysis on the notum. We found that compared to wild-type scales, the upper part of the scales (23.718±0.960 µm) was significantly (t=-20.799, df=6, p<<0.0001) narrower in *Aa-forked* mutants (2.87±0.287 µm, Table 2). On the other hand, in *Aa-forked* mutants the scales were significantly longer then WT (78.75±4.278 µm and 56.789±3.891 µm, respectively. Table 2).

**Table 2.**
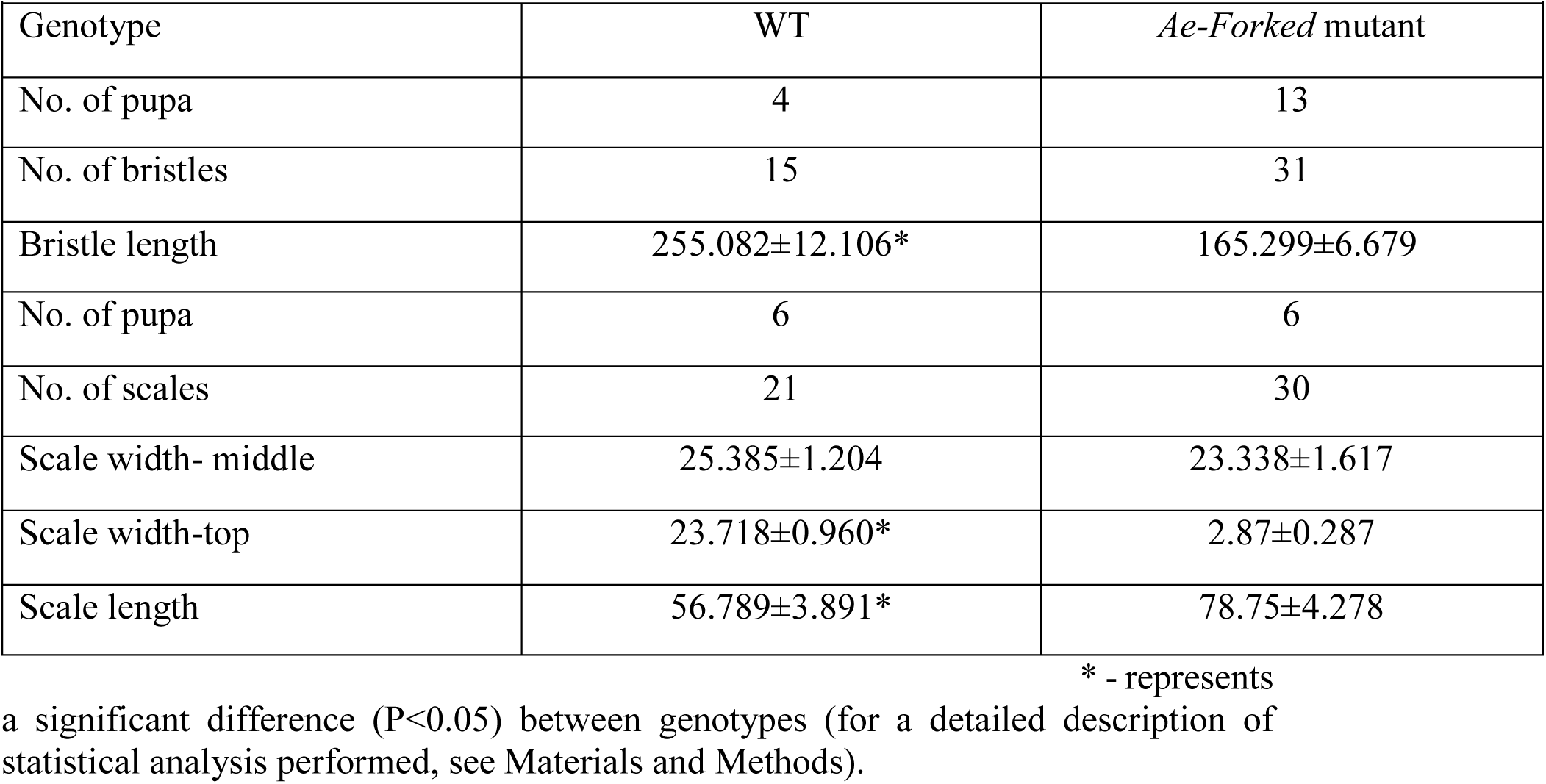
Bristle and scale measurement in WT and *Ae-Forked* mutant.

**Figure 6.**
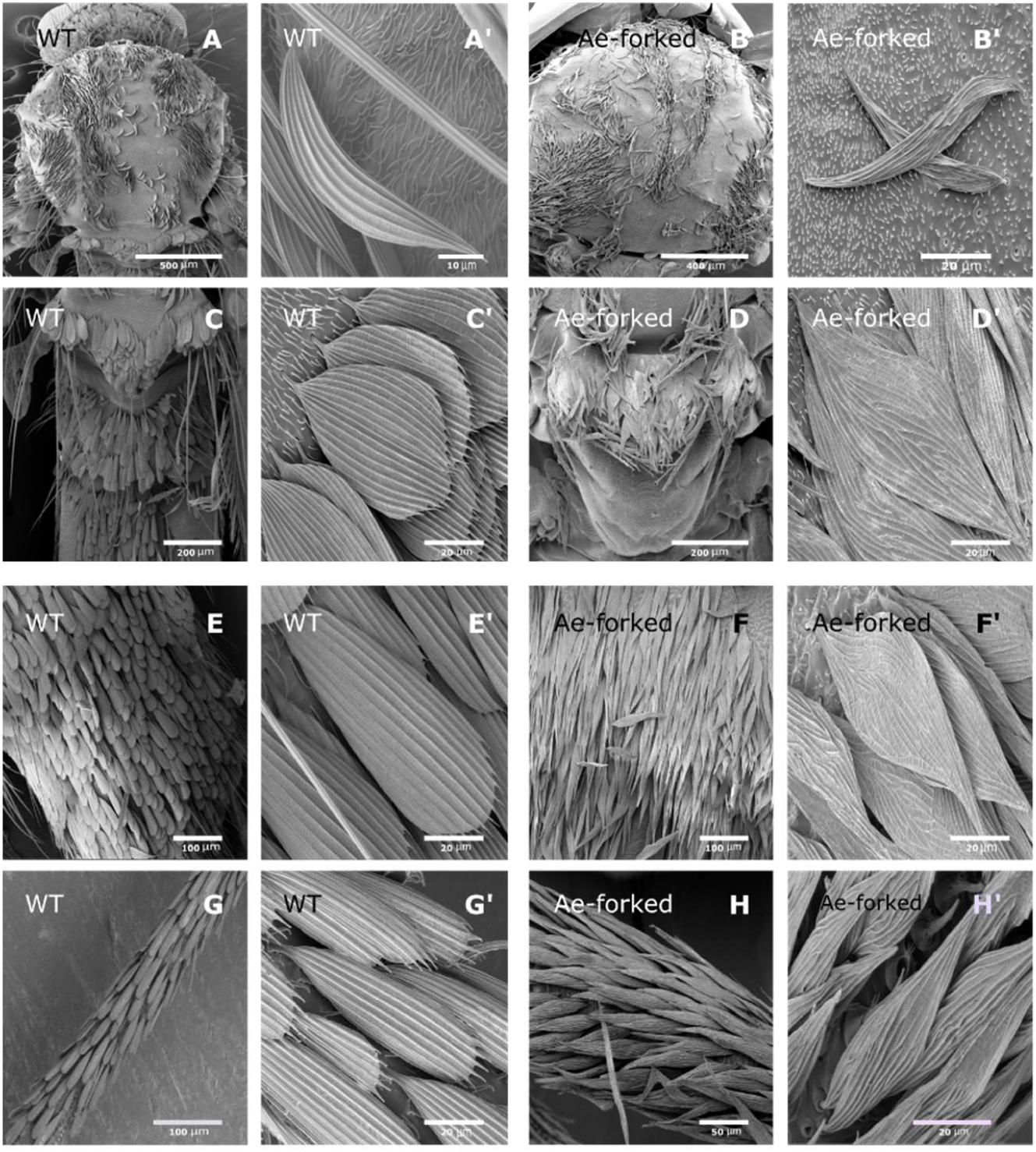
Scanning electron microscopy (SEM) images of scale from different tissues of WT and *Ae-forked* mutnats mosquito *Aedes aegypti*. Thorax of WT (A) and *Ae-forked* (B) mutnats mosquito. Scutellum of WT (C) and *Ae-forked* mutnats (D) mosquito. Abdomen of WT (E) and *Ae-forked* (F) mutnats mosquito. Legs of WT (G) and *Ae-forked* (H) mutnats mosquito. (A’,C’, E’ and G’) high-magnification image of WT scales reveals that along each scale, there are longitudinal ridges which spread from the base of the scale to the tip. High-magnification image of scales from all tissues (B’,D’, F’ and H’) revealed that the ridges on the scales are disorientated. In all spatulate scales (D’, F’ and H’), the upper part was no longer broad, but instead tapered towards the tip, resembling the bristle tip. Scale bars are found on each image.

Since in *Drosophila* the *forked* gene affects both bristle and hair development, we tested whether these structures were affected in in *Aa-forked* mutants. Analyzing bristle morphology in *Aa-forked* mutants (Fig. 7B) revealed that bristles were significant shorter (t=-6.494, df=5, p=0.0013) in *Aa-forked* mutants (165.299±6.679) compared to wild-type (255.082±12.106 µm, Table 2). Closer examination revealed that these bristles lost their regular ridge and valleys pattern (Fig. 7A’ vs. Fig. 7B’).

**Figure 7.**
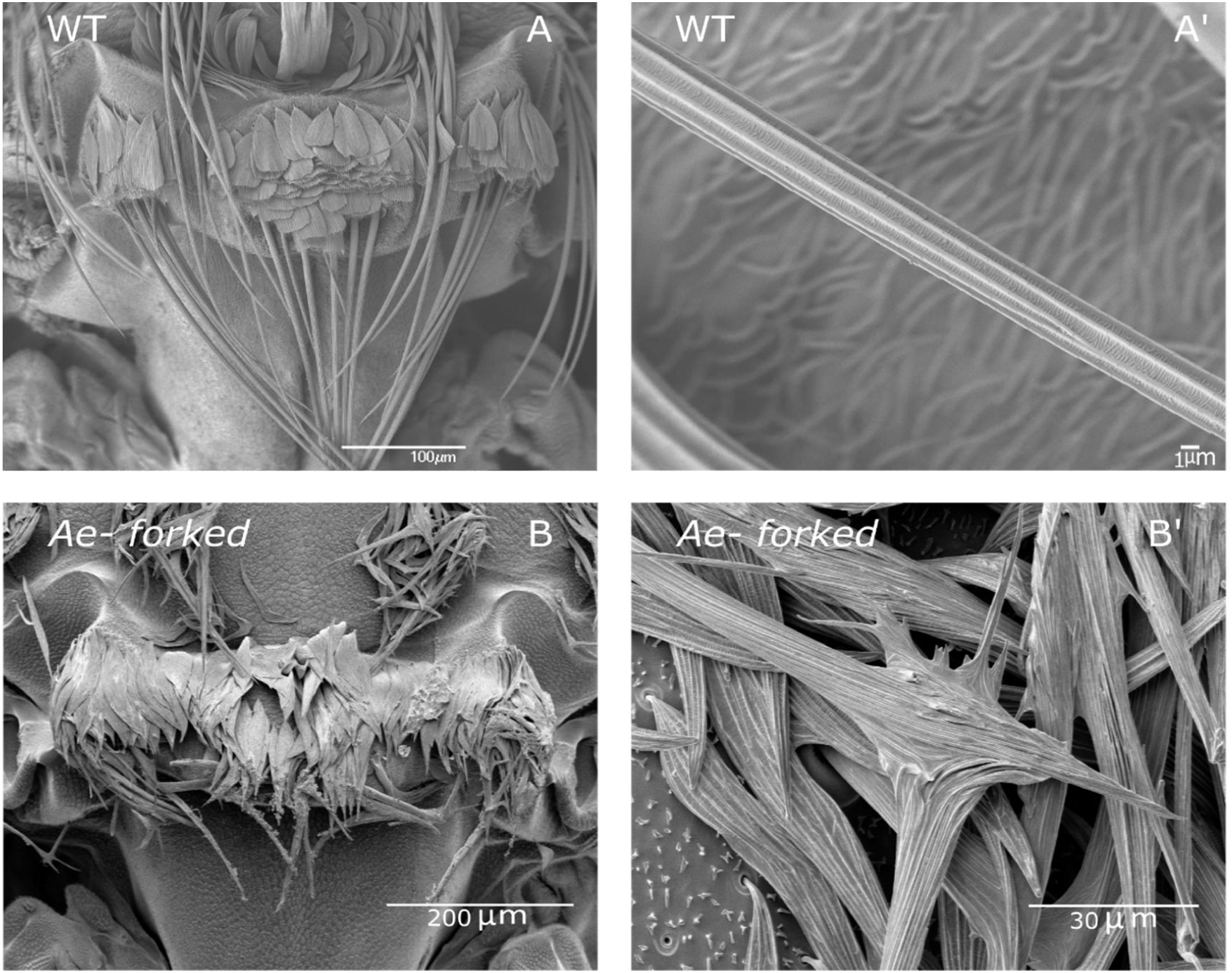
Scanning electron microscopy (SEM) images of bristle from different tissues of WT and *Ae-forked* mutnats mosquito *Aedes aegypti*. Scutellum of WT (A) and *Ae-forked* mutnats (B) mosquito. In *Ae-forked* mutnats, bristle are much shoretr than WT. (A’, B’) high-magnification image of WT (A’) and (B’) *Ae-forked* mutnats bristle. In WT, there are longitudinal ridges from the base of the bristle to the tip. In *Ae-forked* mutnats, the bristle is splitted and the risges are mis-orienated. Scale bars are found on each image.

Next, we analyzed hair structure on the thorax (Fig, 8A-B) and notum (Fig. 8C-D) and found that hair morphology in *Aa-forked* was affected with splintered ends compared to the tapered cylinder morphology of the wild-type; (Fig. 8 A and C vs Fig. 8 B and D). Similar to bristles, the length of hairs were significantly shorter in *Ae-forked* mutant both on the thorax (wild-type −4.289±0.433 µm, mutant - 2.135±0.127 µm, t=-4.776, df=2, p=0.041) and on the notum (wild-type 5.836±0.102 µm, 2.474±0.183 µm, t=-16.061, df=6, p<<0.0001).

**Figure 8.**
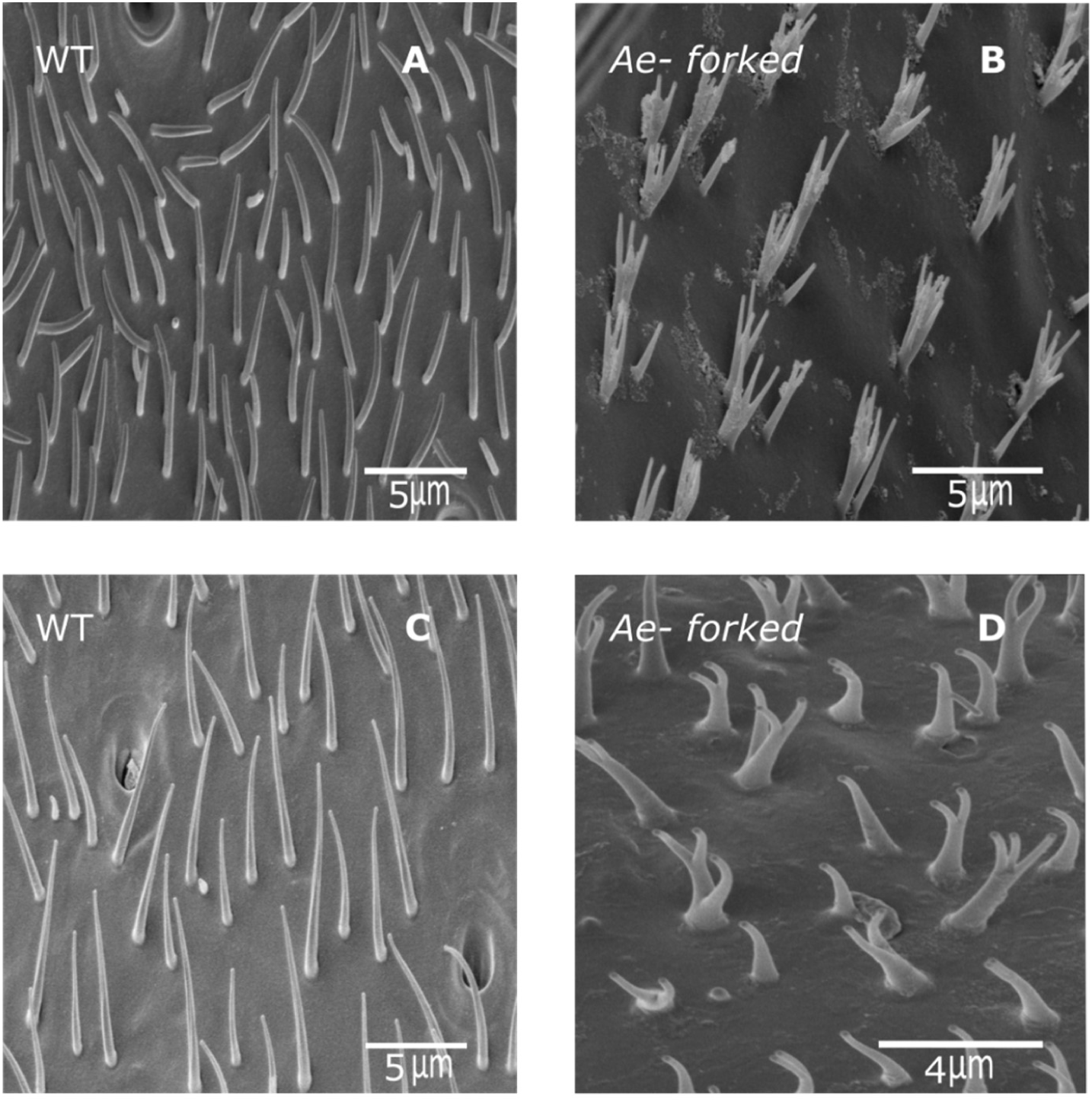
Scanning electron microscopy (SEM) images of hair from different tissues of WT and *Ae-forked* mutnats mosquito *Aedes aegypti*. Hair from the thorax (A,B) and Scutellum (C,D) from mosquito tissues. WT (A,C) and *Ae-forked* mutnats (B, D) mosquito. In both WT tissue, hair are cylinder-like in shape which tappered towrds the tip. In *Ae-forked* mutnats, hair are shorter and split. Scale bars are found on each image.

## Discussion

### Mosquito scale undergoes drastic morphology and actin cytoskeleton changes **d**uring development

In this study we structurally characterized *Aedes aegypti* scale development. We showed that at early developmental stages, scales (like bristles), are cylindrical in shape, with membrane associated actin bundles. Thus, bristles and scales share similar cytoskeleton organization. However, during late scale development, scales undergo drastic morphological changes, which appears in two stages. First, the cylindrical shape changes to a flatter shape, and then second, longer membrane invagination (presumably the future scales ridges) appears on one side of the scale. In bristles, on the other hand, one obvious morphological change occurs, the appearance of ridges and valleys. This change corresponds to the appearance of longer membrane invagination in scales. In both cases, the ridges or longer membrane invaginations correspond to sites on the cell membrane where actin bundles are absent.

The organization of actin bundles also alter during development. Three main phases in actin bundle formation can be described. At early stages, almost the entire circumference of the scale is covered with approximately 20 actin bundles. At the second phase, the actin bundles grow in size and are now triangular in shape as bristle actin bundles. At the third phase, the actin bundles are organized asymmetrically, where one side contains large rounded actin bundles, and at the opposite side, the actin bundles are smaller and flatter. The same kind of asymmetrical actin bundles organization could be found also in bristles, where they are large in cross-sectional area on the inferior surface and small or in some cases absent on the superior surface (Tilney et al., 1995). Whereas the first and second phase of actin bundles organization is found at cylinder shape of the scale, the third phase is associated with the flat shape of the scale with the longer membrane invagination. Thus, although actin bundle organization between scales and bristles showed some similarity, we believe that the different organization of the actin cytoskeleton between these two homologous structures dictates their different morphology.

### Temporal and spatial scale organization

In this study, we revealed that the timing of scale initiation between different body parts (scutellum and legs) is different. On the legs, scale buds occur early on at 5 hours APF compared to scales on the scutellum, which initiate at 7 hours APF. For our experimental purpose, it was important to reveal the exact timing of bud initiation so we could interfere with scale growth early on. Similar phenomena was observed on butterfly wings, where it was shown that longer scales will bud earlier and elongate more quickly than shorter scales (Dinwiddie, et al 2014). In parallel to their different bud initiation time, we also noticed that the pattern of scale development between these two body parts is different. On the legs but not on the scutellum, we noticed that scales occur in cluster of 4 cells that differed in their size. The *Drosophila* mechanoreceptor bristle complex comprises 4 specialized cells (the bristle shaft, socket around its base, bipolar neuron, and neuron sheath). All of these cells derive from a single sensory organ precursor (SOP) cell, which undergoes two asymmetric divisions with subsequent differentiation of the daughter cells (reviewed in Schweisguth, 2015). Since scales lack neuron, and neuron sheath, but do possess socket cell (Greenstein, 1972; Gallant et al., 1998), our finding that on the legs scales and their socket cells occur in cluster of 4, suggests that this cluster of cells may originate from single sensory organ precursor (SOP) cell, which undergoes through 3 division but not 2 as in *Drosophila* SOP lineage. Our results revealed that the pattern of scale development is different between different mosquitoo body parts, suggesting that there are multiple mechanisms that control mosquito scale development.

### In contrast to mosquito bristle, hair and butterfly scales, actin bundles in mosquito play different role in scale elongation

One of the main requirements of actin bundles in *Drosophila* bristles and hairs and in butterfly scales is elongating cell extension. In *Drosophila*, treatment with CD before and after bristle initiation affects bristle elongation (Turner and Adler, 1998; Tilney et al., 2000a). In the butterfly *Vanessa cardui*, inhibition of actin bundle formation before and after bud initiation reveals that actin bundles are required for initial scale elongation (Dinwiddie, et al 2014). In this study, we inhibited actin bundle formation before scale and bristle initiation and found that this treatment affects scale and bristle morphology. As expected, the regular ridge pattern of both cell types was strongly affected. But, whereas bristle was significantly shorter than control, scale were significantly narrower and longer than control scales.

These results led us to further investigate the role of actin bundles in shaping the mosquito’s epidermal cellular extension by mutating one of the known actin bundles proteins, forked. *Drosophila forked* is require for actin bundle formation both for bristles (Tilney et al., 1995) and hairs (Guild et al., 2005; Dickinson and Thatcher, 1997). In *Drosophila forked* mutants, bristles are shorter (50% as the long as wild-type) and thicker, twisted and in some cases exhibit forked tips (Guild et al., 2003; Tilney et al., 2004). We knocked out the mosquito *forked* homologue, *Ae-forked*, and found that, similarly to *Drosophila* bristles, the bristles were significantly shorter than wild-type with mis-organized longitudinal ridges. On the other hand, in *Ae-forked* mutant, scales were significantly longer than wild-type, and lost their ridge pattern. The most affected region in the scales was the tip region. In all three body parts examined, the broad distal region become tapered as bristle tip. In total, our results (both inhibition of actin polymerization and also from *Ae-forked* mutant) demonstrate that actin bundles play differential roles in scale compared to bristles. In bristles, actin bundles are required for cell elongation and in scales actin bundles are required for width formation.

What is the difference in actin bundles organization between scales and bristle that affect their function? We found that the density of ∼20 membrane associated actin bundles is higher in scales compared to bristle. First, both in macrocheta (reviewed in Tilney and DeRosier, 2005), and in scales (this study), actin bundles are evenly distributed along the cell circumference. However, whereas the distance between each actin bundles is about ∼1 µm in macrocheta, in scales, before cells flatten, the distance between each actin bundle is about ∼0.15 µm. Thus, reducing the number of membrane-associated actin bundles and mis-orienting them by inhibiting actin polymerization chemically or by gene deletion led to narrower scales, with strong effects on scale tip widening. Thus, the unique dense membrane-associated actin bundles in scales may contribute for widening of the scale.

## Materials and methods

### *Aedes aegypti* Mosquito Rearing

Adult *A. aegypti* mosquitoes were reared in a growth chamber set at 27 °C and 75% humidity, on a 12 h light: 12 h dark cycle, with unlimited access to water and 10% sucrose solution on a cotton wick. Larvae were reared at 27 °C and 75% humidity in water and fed with a mixture of Tetramin fish flakes and yeast (1:1 ratio). Female mosquitoes fed on mouse blood using a Hemotek feeder (PS-6 System, Discovery Workshops, Accrington, UK). Blood (3.0 ml) was transferred into the Hemotek blood reservoir unit and the temperature was set to 37°C system.

### Single guide RNAs (sgRNA) synthesis

The following sgRNA 5 ‘GGAGTCCACCCTGAAGCCAT 3 ‘targeting exon 7 in *Ae-forked* gene was used. The primers for sgRNA design were annealed using Phusion High-Fidelity DNA Polymerase. sgRNA templates were transcribed by using T7 polymerase from T7 Megascript kit. RNA transcripts were purified using MEGAclear™ Kit.

### Embryo microinjections

*A. aegypti* mosquitoes expressing Cas9 protein under control of the ubiquitin L40 (AAEL006511) promoter were used (Li et al., 2017). Embryo microinjections were performed as previously described (Jesinskiene et al, 2007). Briefly, four days after blood feeding, 10 mated gravid females were transferred into plastic tubes with oviposition substrate, and kept in the dark for oviposition. After 30 minutes, eggs were collected and aligned on a piece of Whatman filter paper. Eggs were covered with a 1:1 mix of Halocarbon 700 oil: Halocarbon 27 oil to prevent desiccation. Quartz needles were pulled using a Sutter P2000 needle puller and were used with a Femtojet injector (Eppendorf) and InjectMan micromanipulator. 0.2 nL sgRNA at 400 ng/µL was injected into embryos. After injections, embryos were allowed to recover under insectary conditions for 5 days before hatching them under vacuum into water.

### Molecular analysis of mutant individuals

G_0_ adults were visually screened for bristle and scale defects. Individual legs from putative mosaic and non-mosaic edited G_0_ mutants or G_1_ and G_2_ mosquitoes were collected and genomic DNA extracted using Wizard® Genomic DNA Purification Kits or Qiagen DNeasy® Blood & Tissue kits. Genomic DNA was used as template for PCR with the following primers: Forward 5 ‘CTGTGGGACCCCCACCGCCA 3’ and reverse 5 ‘CTGATATAATGGACATGCTT 3’. PCR amplicons were directly sequenced.

### Pupa phalloidin and antibody staining

For examination of bristles and scales, 0-1-hour old pupa were collected and reared individually. At the appropriate developmental time after pupal formation (as mentioned in the text), pupal cuticles were removed and the pupae were fixed for confocal or electron microscopy. For confocal microscopy pupa were fixed in 4% paraformaldehyde in PBS for overnight. The samples were washed three times with 0.3% Triton X-100 in PBS for 10 min. each time. For phalloidin staining, samples were washed three times with 0.3% Triton X-100 in PBS for 10 min each time and then incubated overnight with phalloidin. Then, the samples were washed three times in 0.3% Triton X-100 in PBS. For antibody staining, the thoraces were blocked in 0.1% Triton X-100 containing 4% bovine serum albumin for 1 h. The samples were then incubated overnight with a primary antibody in the blocking solution at 4°C, washed three times in 0.3% Triton X-100 in PBS, and incubated with secondary antibodies in blocking solution for 2 h at room temperature or at 4°C overnight in the dark. After incubation, samples were washed three times with 0.3% Triton X-100 in PBS for 10 min each time. For confocal observation, both phalloidin and antibody-stained samples were placed on a slide and mounted in 50% glycerol. A coverslip was placed on the sample, and the preparation was sealed with nail polish. The slides were examined with an Olympus FV1000 laser-scanning confocal microscope. Mouse anti-α-tubulin (1:250) (Sigma) primary antibodies were used. Goat anti-mouse Cy2 and Cy3 and goat anti-rabbit Cy3 (Jackson ImmunoResearch) secondary antibodies were used at a dilution of 1:100. The goat anti-rabbit Cy3 (Molecular Probes) secondary antibodies were used at a dilution of 1:500. For actin staining, we used Oregon Green 488- or Alexa Fluor 568-conjugated phalloidin (1:250) (Molecular Probes).

### Injection of inhibitors

Stock solution of 5 mg/mL cytochalasin D in DMSO was prepared. Twenty *A. aegypti* pupa at the appropriate developmental time after pupal formation (as mentioned in the text), were injected with 2 nL of cytochalasin-D into the first or second abdominal segment. Control pupae were injected with DMSO. Pupae were allowed to develop and then were prepared for SEM analysis. Each experiment was repeated three times.

### Scanning electron microscopy (SEM)

Samples were fixed and dehydrated by immersion in increasing concentrations of ethanol (30%, 50%, 75%, and twice in 100%; 10 min each). The samples were then completely dehydrated using increasing concentrations of hexamethyldisilazane (HMDS) in ethanol (50%, 75%, and twice in 100%; 2 h each). The samples were air dried overnight, placed on stubs, and coated with gold. The specimens were examined with a scanning electron microscope (SEM; JEOL model JSM-5610LV). Length measurements of adult bristles were performed using Image J (http://rsb.info.nih.gov/ij/) (version 1.40j) software.

### Bristles and scales measurements and statistics

Bristles and scales were examined by scanning electron microscope (SEM). Length and width of bristles and scales were measured from SEM images by using the Image J. The length of the bristles were measured from the base of the bristle (only in cases where we could detect the socket cells on the epidermis) till the bristle tip. In both cytochalasin-D-treated and forked mutants mosquitoes with split bristles, the measurement was restricted to the main bristle shaft only. The length of the scale was measured from the attenuation at the base till the distal part. The width of the scale was measured both at the middle part of the scale and also at the tip part of the scale. For each individual we measured 3-6 different scales or bristle (see Tables for the exact number) and used the average measurement for the statistical analysis. The difference between wild type and treated individuals (either cytochalasin-D or *Ae-forked* CRISPR mutants) was than analysed using t-test with unequal variance estimates.

### Transmission Electron microscope (TEM)

Legs from each developmental stage pupa were dissected as described above and fixed for 20 min in 2% glutaraldehyde in 0.2 M PO_4_ (pH 6.8) at room temperature and then for 1 h on ice. After 1 h, the specimens were transferred to a fixative comprising cold water, 0.2 M PO_4_ (pH 6.2), 4% osmium tetroxide (OsO_4_), and 50% glutaraldehyde and placed on ice for 1 h. The specimens were then washed three times in cold water (20 min each) and incubated in 1% uranyl acetate overnight at 4°C. This was followed by transfer to a dehydration series spanning from 30 to 100% acetone, with 10% increases being made at each 15-min interval. Samples were then treated twice in propylene oxide for 15 min and soaked for 1 h in a 1:1 solution of propylene oxide and araldite, followed by overnight incubation at 4°C in a 1:2 propylene oxide/araldite mixture. The tissues were then transferred to araldite and incubated for 1 h, placed on araldite blocks (the blocks were polymerized the previous day at 60°C), and embedded in araldite. These were then left at room temperature for 30 min after embedding, at which point the samples were oriented and incubated at 60°C for 24 h. Transverse sections were cut through the thorax using a Leica UltraCut UCT ultra microtome equipped with a diamond knife, stained with uranyl acetate and lead citrate, and then examined with a JEOL JEM-1230 transmission electron microscope operating at 120 kV.

## Acknowledgments

We thank for Omar Akbari for generously providing Cas9-expressing *Aedes* line. We thank Alexander Raikhel for teaching us mosquitos rearing and maintenance. This work was supported by the NIH/NIAID grants R01AI116636, R01AI128201, R01AI150251, and NSF/BIO grant 1645331 to JLR

## References

Bryan, J., Edwards, R., Matsudaira, P., Otto, J. and Wulfkuhle, J. (1993). Fascin, an echinoid actin-bundling protein, is a homolog of the Drosophila singed gene product. Proc. Natl. Acad. Sci. U. S. A. 90, 9115–9119.

Cooper, J. A. (1987). Effects of cytochalasin and phalloidin on actin. J. Cell Biol. 105, 1473–1478.

Day, C. R., Hanly, J. J., Ren, A. and Martin, A. (2019). Sub-micrometer insights into the cytoskeletal dynamics and ultrastructural diversity of butterfly wing scales. Dev. Dyn. 248, 657–670.

Dickinson, W. J. and Thatcher, J. W. (1997). Morphogenesis of denticles and hairs in Drosophila embryos: Involvement of actin-associated proteins that also affect adult structures. Cell Motil. Cytoskeleton 38, 9–21.

Dinwiddie, A., Null, R., Pizzano, M., Chuong, L., Leigh Krup, A., Ee Tan, H. and Patel, N. H. (2014). Dynamics of F-actin prefigure the structure of butterfly wing scales. Dev. Biol. 392, 404–418.

Fei, X., He, B. and Adler, P. N. (2002). The growth of Drosophila bristles and laterals is not restricted to the tip or base. J. Cell Sci. 115, 3797–3806.

Gallant, J. L., Connor, C. E. and Van Essen, D. C. (1998). Neural activity in areas V1, V2 and V4 during free viewing of natural scenes compared to controlled viewing. Neuroreport 9, 1673–1678.

Greenstein, M. E. (1972). The ultrastructure of developing wings in the giant silkmoth, Hyalophora cecropia. II. Scale-forming and socket-forming cells. J. Morphol. 136, 23–51.

Guild, G. M., Connelly patricia S., Vranich, K. A., Shaw, M. K. and Tilney, L. G. (2002). Actin filament turnover removes bundles from Drosophila bristle cells. J. Cell Sci. 115, 641–653.

Guild, G. M., Connelly, P. S., Ruggiero, L., Vranich, K. A. and Tilney, L. G. (2003). Long continuous actin bundles in Drosophila bristles are constructed by overlapping short filaments. J. Cell Biol. 162, 1069–1077.

Guild, G. M., Connelly, P. S., Ruggiero, L., Vranich, K. A. and Tilney, L. G. (2005). Actin filament bundles in Drosophila wing hairs: Hairs and bristles use different strategies for assembly. Mol. Biol. Cell 16, 3620–3631.

Jasinskiene, N., Juhn, J. and James, A. A. (2007). Microinjection of A. aegypti embryos to obtain transgenic mosquitoes. J. Vis. Exp.

Li, M., Bui, M., Yang, T., Bowman, C. S., White, B. J. and Akbari, O. S. (2017). Germline Cas9 expression yields highly efficient genome engineering in a major worldwide disease vector, Aedes aegypti. Proc. Natl. Acad. Sci. U. S. A. 114, E10540–E10549.

Mitchell, H. K., Roach, J. and Petersen, N. S. (1983). The morphogenesis of cell hairs on Drosophila wings. Dev. Biol. 95, 387–398.

Paterson, J. and O’Hare, K. (1991). Structure and transcription of the singed locus of Drosophila melanogaster. Genetics 129, 1073–1084.

Ren, N., He, B., Stone, D., Kirakodu, S. and Adler, P. N. (2006). The shavenoid gene of Drosophila encodes a novel actin cytoskeleton interacting protein that promotes wing hair morphogenesis. Genetics 172, 1643–1653.

Schweisguth, F. (2015). Asymmetric cell division in the Drosophila bristle lineage: From the polarization of sensory organ precursor cells to Notch-mediated binary fate decision. Wiley Interdiscip. Rev. Dev. Biol. 4, 299–309.

Tilney, L. G. and DeRosier, D. J. (2005). How to make a curved Drosophila bristle using straight actin bundles. Proc. Natl. Acad. Sci. U. S. A. 102, 18785–18792.

Tilney, L. G., Tilney, M. S. and Guild, G. M. (1995). F actin bundles in Drosophila bristles I. Two filament cross-links are involved in bundling. J. Cell Biol. 130, 629–638.

Tilney, L. G., Connelly, P., Smith, S. and Guild, G. M. (1996). F-actin bundles in Drosophila bristles are assembled from modules composed of short filaments. J. Cell Biol. 135, 1291–1308.

Tilney, L. G., Connelly, P. S., Vranich, K. A., Shaw, M. K. and Guild, G. M. (2000a). Actin filaments and microtubules play different roles during bristle elongation in Drosophila. J. Cell Sci. 113, 1255–1265.

Tilney, L. G., Connelly, P. S., Vranich, K. A., Shaw, M. K. and Guild, G. M. (2000b). Regulation of actin filament cross-linking and bundle shape in Drosophila bristles. J. Cell Biol. 148, 87–99.

Tilney, L. G., Connelly, P. S., Ruggiero, L., Vranich, K. A., Guild, G. M. and DeRosier, D. (2004). The role actin filaments play in providing the characteristic curved form of Drosophila bristles. Mol. Biol. Cell 15, 5481–5491.

Turner, C. M. and Adler, P. N. (1998). Distinct roles for the actin and microtubule cytoskeletons in the morphogenesis of epidermal hairs during wing development in Drosophila. Mech. Dev. 70, 181–192.

Wu, C. W., Kong, X. Q. and Wu, D. (2007). Micronanostructures of the scales on a mosquito’s legs and their role in weight support. Phys. Rev. E - Stat. Nonlinear, Soft Matter Phys. 76,.

Zhou, B., Williams, D. W., Altman, J., Riddiford, L. M. and Truman, J. W. (2009). Temporal patterns of broad isoform expression during the development of neuronal lineages in Drosophila. Neural Dev. 4,.

